# N-Cadherin mediated cell rearrangements shape embryonic macrophage cluster

**DOI:** 10.1101/2024.06.05.597542

**Authors:** Jacob Hasenauer, Xiang Meng, Honor Scarborough, Jasmine A. Stanley-Ahmed, Darius Vasco Köster, Aparna Ratheesh

**Author notes:** Contributed equally (Jacob Hasenauer, Xiang Meng).

## Abstract

Drosophila embryonic macrophages are highly motile phagocytic and secretory cells which are essential for embryonic development. Embryonic migration and dispersal of these cells along pre-determined routes is invariably tied to their functions such that misrouting or delay in macrophage migration has serious consequences for embryogenesis and adult homeostasis. In this study, we describe early steps of macrophage migration from their site of origin in the head mesoderm to the germband and show that hemocytes start moving as an epithelial-like cluster with N-Cadherin based Adherens junctions. The cells within the cluster move in a co-ordinated manner and exhibit cell rearrangements mediated through junction shrinkage. We found that N-Cadherin is dynamically relocalized during cluster extension and modulating N-Cadherin levels results in hemocyte dispersal away from their main axis of migration which affects migration into the germband. We therefore elucidate a novel multi-tiered mechanism which ensures that macrophages are positioned appropriately to regulate their distribution in the embryo.

## Introduction

*Drosophila* embryonic hemocytes are highly migratory cells which bear striking similarities to their mammalian macrophage counterparts in migratory behaviour, function and ontogeny^1–4^. During embryonic development, hemocytes disseminate throughout the embryo following seemingly programmed developmental pathways^5–8^ following cues from the platelet-derived growth factor- and vascular endothelial growth factor-related factors (Pvf) 2 and 3 ^6,8–10^. During their initial dispersal and further movement during the embryonic life cycle, they phagocytose dead cells and secrete ECM components essential for embryonic development^1,11–13^.

Much of the information on hemocyte migratory mechanics come from studies on their migration along the ventral surface of the embryo beneath the epithelium at late stages of embryogenesis (stage 12 and later). During these stages, they migrate as mesenchymal single cells and exhibit microtubule dependent contact inhibition of locomotion which is essential for efficient embryonic dispersal ^14–16^. They exhibit large actin and microtubule rich lamellipodia capable of force generation and transduction which is dependent on the activity of the small GTPases such as Rac and Rap1^14,17–21^. However not much is known about the migration dynamics of hemocytes eat early stages of delamination and initiation of dispersal. In this study, we unravel the molecular changes that occur within hemocytes to allow delamination and migration to the germband which sets the stage for hemocyte dispersal across the embryo.

## Results & Discussion

Hemocytes are specified in a sharply delimited area within the procephalic mesoderm on the ventral side of the embryo at the blastoderm stage^5^. By early stage 11 of development, the first hemocytes reach the extended germband which it infiltrates^20,22^. To understand hemocyte migration dynamics before they reach the germband, we used two-photon microscopy to image hemocytes in embryos from stage 8 onwards when we could first detect hemocyte nuclei under the serpent promoter which specifically labels hemocytes^23^. At stage 8, lateral views of the embryo revealed that the prohemocytes form a cluster in a roughly triangular shape restricted to the ventral side (Fig 1a-b, Video 1). We then imaged these cells for the next 2 hours, focusing on the cells which migrate into the germband. Interestingly, the hemocyte cluster undergoes a remarkable change in shape, elongating along the anterior-posterior axis as more cells join the back of the cluster (Video 1) with the front of the cluster reaching the yolk on the dorsal side by early stage 11 (Fig 1b-c). We analysed changes in cluster parameters in fixed embryos labelling both hemocyte nuclei and Actin; imaging the entire cell population further confirmed that these shape changes are accompanied by an increase in total hemocyte numbers (Fig 1d, Fig S1a). We then analysed cell organization within the cluster in live embryos and found that during the elongation phase, hemocytes displayed prominent cell-cell contacts and moved together as an interconnected cluster (Fig 1e, Video 1). This is surprisingly different to hemocyte behaviour at later stages when they migrate as single cells or in chains forming transient cell-cell contacts ^14,16,18,22^. At stage 8, small protrusions were restricted to a few cells at the front of the cluster facing the germband (Fig 1e, video 1). Protrusions remained restricted to the leading cells at stage 11 as cells started leaving the cluster in chains along the presumptive routes of migration towards the germband (Video 1, Fig S1b).

**Figure 1.**
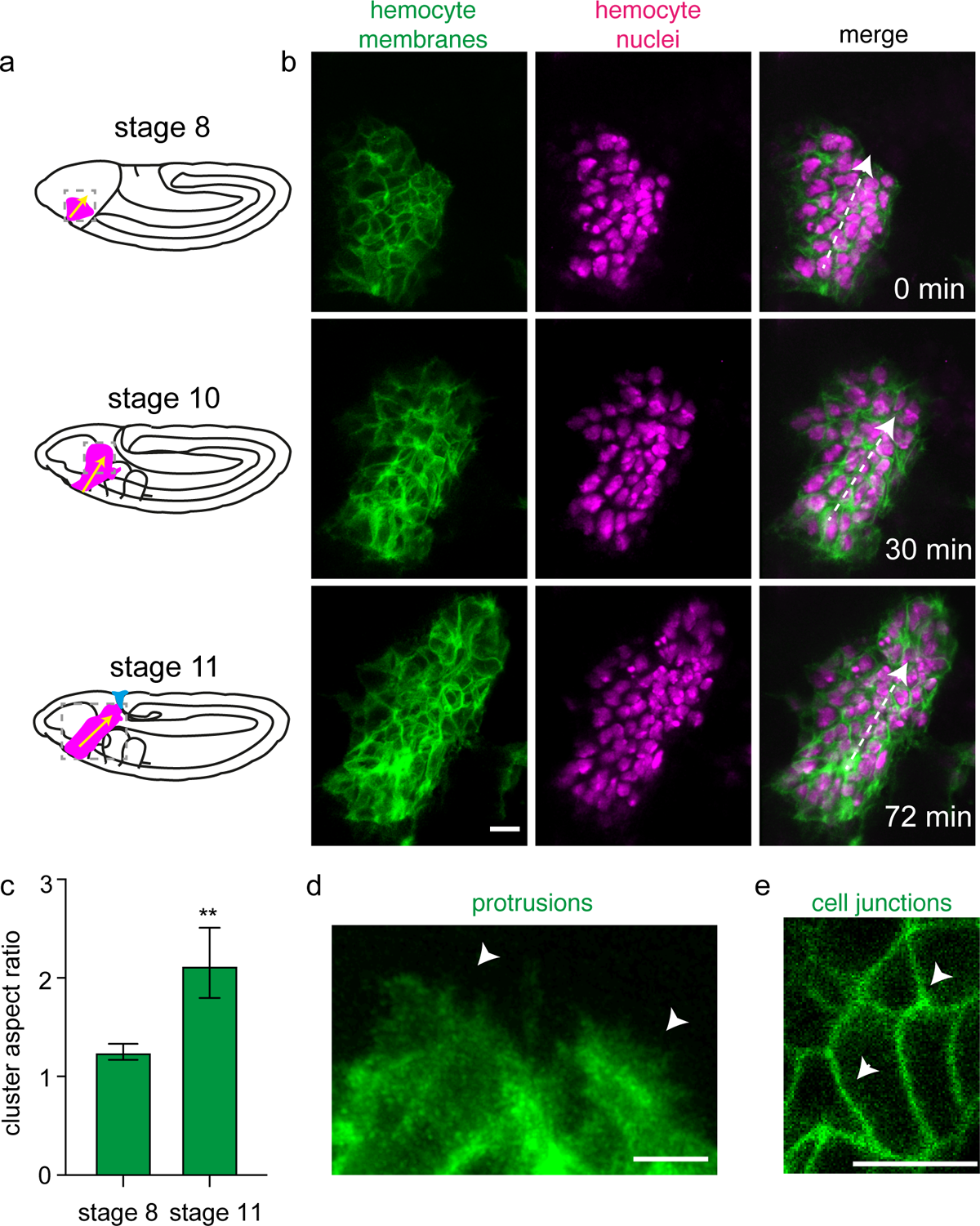
**(a)** Cartoon showing the shape of the hemocyte cluster (Magenta) migrating towards the germband at stage 8, 10, and 11. Yellow arrows indicate the direction of hemocyte migration towards the germband. **(b)** Representative images from 3D projection of multiphoton time-lapse imaging of a *srpHemo>myrGFP; srpHemo-H2A::3xmCherry* embryo from stage 8 (0 mins) to stage 10 (30 mins) and late stage 11 (72 mins), in which *GFP* labels hemocyte cell membranes (green) and *mCherry* labels hemocyte nuclei (magenta), along with a merge of both channels. The position of the cluster in the embryo at different timepoints is indicated by the dotted-line boxes in (A). White arrows indicate the direction of hemocyte migration. Scale bars = 10 µm. **(c)** Quantification of the shape of the hemocyte cluster in stage 8 and stage 11 from live imaging of *srpHemo-H2A::3XmCherry* embryos revealed a significant increase in the aspect ratio of the cluster. n = 3 embryos. **(d)** Representative zoomed in image from 3D projection of multiphoton time-lapse imaging of a *srpHemo>myrGFP; srpHemo-H2A::3xmCherry* embryo at the edge of the hemocyte cluster at 30 minutes from **(b)**. Arrowhead indicates protrusions at the leading edge of the cluster. Scale bar = 5 µm. **(e)** Representative image of a single z slice from multiphoton time-lapse imaging of a *srpHemo>myrGFP; srpHemo-H2A::3xmCherry* embryo in the interior of the hemocyte cluster at 30 minutes. White arrows indicate cell junctions between hemocytes. Scale bar = 10 µm.

We next sought to understand the cell behaviours which underlie cluster elongation. We analysed the migratory behaviour of the cells in the cluster in bulk by imaging hemocyte nuclei with high spatiotemporal resolution (every 40 seconds) for about an hour (Video 2). We detected individual nuclei by 3D segmentation (Fig S2a) and plotted the nuclear position over time. At early stage 8, the cells of the cluster were distributed fairly evenly around the centroid of the cluster (Fig 2a) and by stage 11, the cells reorganized themselves along the axis of migration (Fig 2b-d, Fig S2b) suggesting that active cell movements underlie the cluster’s elongation. We then tracked individual nuclei of the cluster manually and characterized migration parameters. Hemocytes undergo 4 rounds of cell division between stages 6 and 11^5^ and we observed division events during this period (Fig 2e, video 2), leading us to conclude that proliferation contributes towards increase in cell number during cluster elongation. To analyse cell migration independent of division events, we omitted division events manually from all our further analysis. The cell movements within the cluster showed average speeds of about 2.6 microns per minute (Fig 2f). We then asked whether the cell movements within the cluster are correlated with the direction of migration of the cluster itself. Correlation analysis of 3D velocity vectors showed that the average spatial correlation was positive over the period of elongation (Fig 2g). Frequency analysis of movement correlation showed that negatively correlated movements did occur within the cluster, but a higher fraction of cells displayed positive correlations (Fig 2h) suggesting that, on the whole the cells within the cluster moved with high spatial correlation.

**Figure 2.**
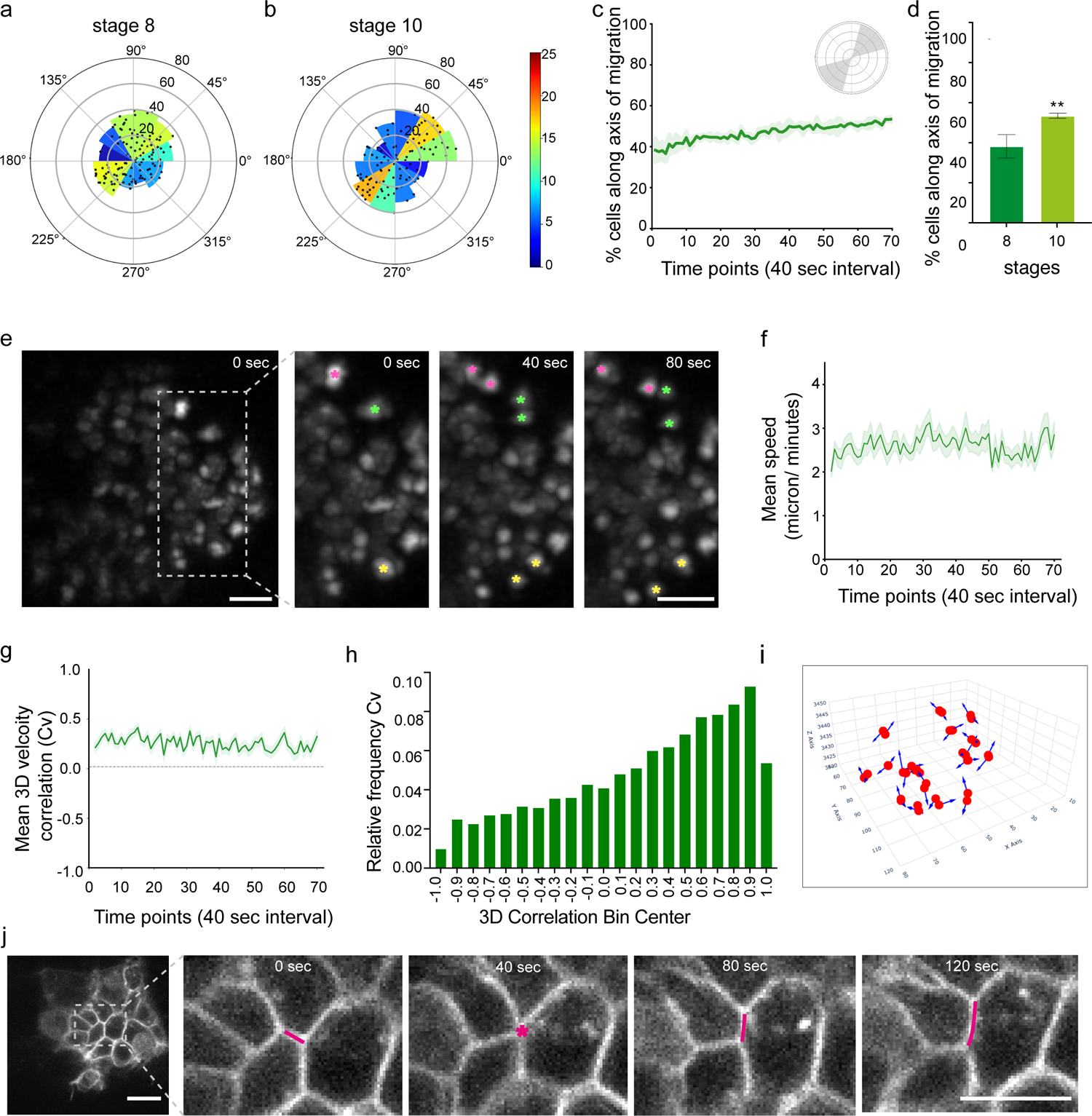
**(a, b)** Polar histogram illustrating the spatial distribution of nuclei (control) between stage 8 (0 minutes) and stage 10 (46 minutes) from live imaging of *srpHemo-H2A::3XmCherry* embryos. Each dot represents the centroids of nuclei segments. The radial axis indicates the physical distance of the displacement in microns, and the angular axis represents the direction in degrees. Colour scale on the right indicates the nucleus number in each 30-degree angle range (histogram bin=12). **(c)** Line graph showing the percentage of cells in the range of polar histogram from 15-75 degree (1st quadrant, grey shade area in cartoon and Fig S2) and 195-255 degree (3rd quadrant, grey shade area in cartoon) over time from stage 8 (0 minutes) and stage 10 (46 minutes) representing the cells along the main axis of migration. The shaded area around the line represents the standard error of the mean (SEM). **(d)** Bar graph comparing the percentage of cells in (grey shade area in cartoon in **c**) at two specific time points, stage 8 (0 minutes) and stage 10 (46 minutes). Error bars denote the standard error of the mean (SEM). Statistical significance between t0 and t70 is indicated with ** (p < 0.01). **(e)** Representative images from multiphoton time-lapse imaging of *srpHemo-H2A::3xmCherry* embryo showing cell division events over subsequent time points separated by 40 seconds. The coloured asterisks denote dividing cells. Scale bar = 10 µm. **(f, h)** Line graphs showing the mean speed **(f)** and 3D velocity cluster correlation **(h)** of hemocytes in *srpHemo-H2A::3xmCherry* embryos over time from stage 8 (0 minutes) and stage 10 (46 minutes). The shaded area around the line represents the standard error of the mean (SEM). **(g)** Relative frequency of 3D velocity cluster correlation from individual cells over the entire 46 minutes. Representative 3D plot from manual tracking of srpHemo-H2A::3xmCherry embryo showing the spatial direction of displacement of dividing cells (red) denoted by arrows and tracks (blue). **(j)** Representative images from multiphoton time-lapse imaging of a *srpHemo>myrGFP; srpHemo-H2A::3xmCherry* embryo showing T1 transitions indicating junction shrinkage and cell rearrangements (magenta asterisk and lines). Scale bar=5 microns.

We then tried to unravel the mechanisms of cluster elongation. Since the shape change of the cluster was highly reminiscent of convergent extension (CE) movements during gastrulation^24–27^, we turned to cell behaviours underlying CE. Oriented cell divisions play a role in CE movements^28,29^, however the directional displacement of hemocytes post divisions was seemingly random (Fig 2i, Video 4) suggesting that while division events increase the number of cells within the collective, oriented cell divisions do not contribute to cluster elongation. We then analysed whether neighbour exchange events, also called intercalation, occur during cluster elongation. We imaged the hemocyte membrane as well as nucleus and found that indeed, cells within the cluster exchange neighbours (Video 5). Interestingly, intercalation events seemed to occur through T1 transitions leading to junction shrinkage (Figure 2j) and neighbour exchange similar to those described during epithelial CE movements ^24,25,30,31^. The intercalation events occurred scattered throughout the cluster apparently stochastically.

Since cells within the cluster displayed cell-cell contacts, we then checked whether the cluster is epithelial in nature and looked at classical Adherens junction components. We first assessed E-Cadherin within the cluster and found that at stage 8, hemocytes displayed E-Cadherin at the contacts, but this was highly transient and was completely absent within the cluster by stage 10 (Fig S3a-b). Interestingly, we found that N-Cadherin was expressed at cell junctions and the cells displayed beta catenin and an circumferential actin belt until the end of stage 11 when hemocytes started leaving the cluster in chains when the N-Cadherin appeared more cytoplasmic (Fig 3a-c,e, Fig S3c). Live imaging of N-Cadherin using a N-Cadherin knock in eGFP line^32^ showed that N-Cadherin appeared as highly dynamic puncta (Fig3d, Video 6). At stage 8, the junctional intensity of N-Cadherin was not significantly different between the cells at the outer edges of the cluster and the interior (Fig 3e). However, as the cluster elongation progressed, N-Cadherin increased at cell junctions, particularly at those junctions in the inner region of the cluster (Fig 3d). Cell density heatmaps within the cluster indicated a similar result with the density being highest in the middle of the cluster where cells are tightly packed with lower area and shorter junctions (Video 7, Fig 3f-g). Our data suggests that hemocyte cluster contains a core of cells which move as interconnected epithelial cells while the cells at the front appear more mesenchymal.

**Figure 3.**
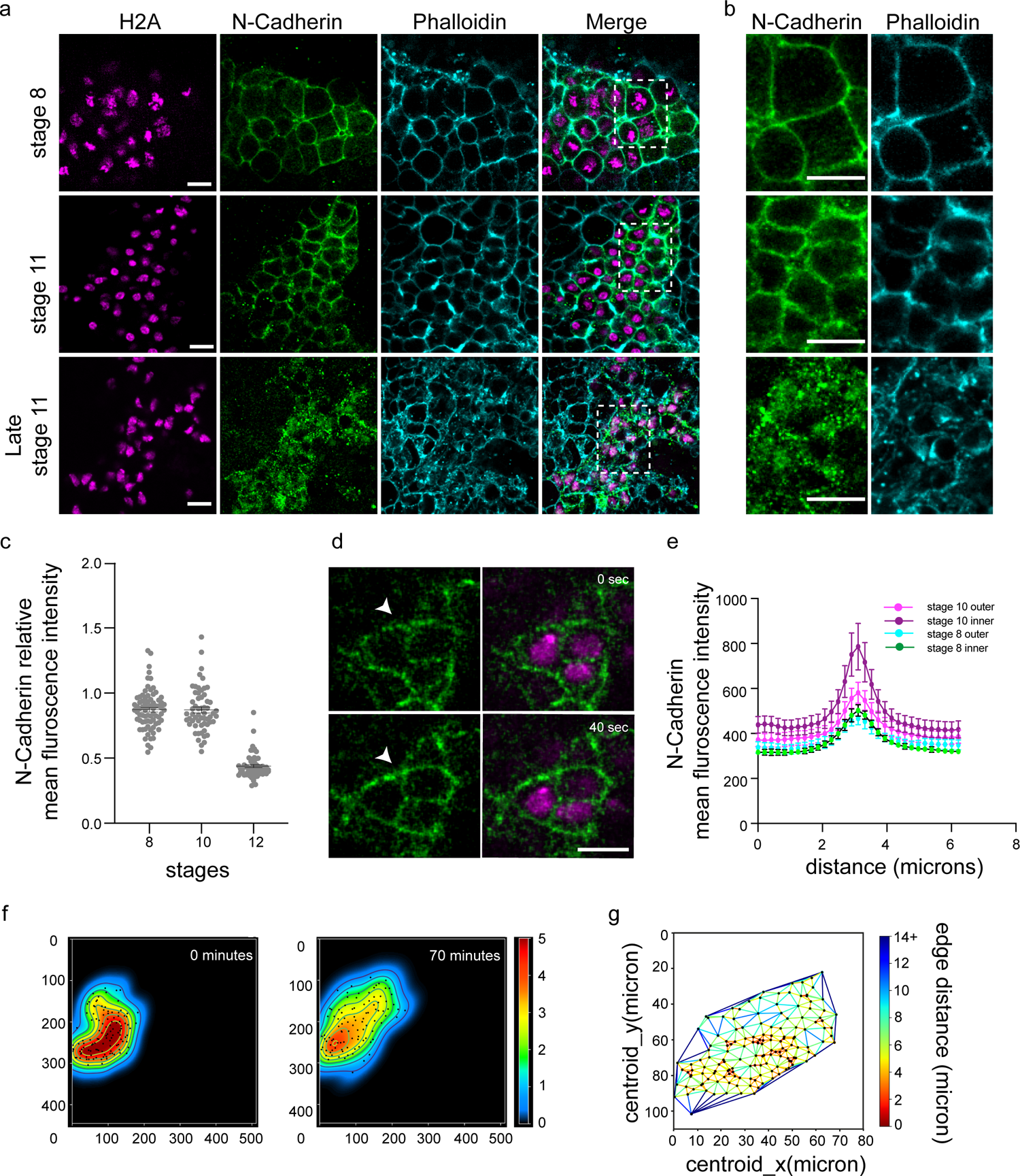
**(a)** Representative images showing a single z slice of fixed confocal microscopy images of *N-Cadherin::GFP; srpHemo-H2A::3XmCherry* embryos across stage 8, 10, and late stage 11 with *mCherry* labelling hemocyte nuclei in magenta, GFP labelling N-Cadherin in green and Phalloidin in cyan. Scale bars = 10 µm. **(b)** Representative image showing GFP labelled N-Cadherin in green and Phalloidin in cyan within a region of interest as indicated by white dotted boxes from **(a)**, images taken from the same z slices and embryos as presented in **(a).** Scale bars = 10 µm. **(c)** Quantification of *N-Cadherin::GFP* mean fluorescence intensity at hemocyte-hemocyte junctions at stage 8, 10, and 12 in *N-Cadherin::GFP; srpHemo-H2A::3XmCherry* embryos, normalised to the mean fluorescence intensity from *N-Cadherin::GFP* at non-hemocyte mesodermal junctions. n = 12-60 junctions. **(d)** Representative image from multiphoton time-lapse imaging of an *N-Cadherin::GFP; srpHemo-H2A::3XmCherry* embryo from 0 sec to 40 sec. Arrowheads indicate dynamic N-Cadherin puncta. Scale bar = 10 µm. **(e)** Quantification of N-Cadherin junctional intensity by line scan analysis of fluorescence at hemocyte-hemocyte cell contacts in stage 8 and 10 from inner and outer regions in the hemocyte cluster in *N-Cadherin::GFP; srpHemo-H2A::3XmCherry* embryos. Mean intensity profile was plotted with s.e.m.. n = 9-26 junctions. **(f)** Density heatmap utilizing Gaussian KDE to visualize the density of nuclei between from stage 8 (0 minutes) and stage 10 (46 minutes) in *srpHemo-H2A::3xmCherry* embryos. Each dot represents the centroids of nuclei segments. Contour lines indicate areas of increasing density from blue (low density) to red (high density). Colour scale on the right represents density values from 0 (lowest) to 5 (highest). **(g)** Delaunay triangulation illustrating the spatial relationships and connectivity distances between nuclei in a two-dimensional space in *srpHemo-H2A::3xmCherry* embryo. Each dot represents the centroids of nuclei segments. Each edge of the triangulation is colour-coded according to the length in microns. The colour scale on the right represents the edge distances, ranging from 0 microns (blue) to 14 (or >14) microns (red).

We next assessed the role of N-Cadherin in hemocyte cluster elongation. We reduced N-Cadherin levels in hemocytes specifically using an RNAi for N-Cadherin under the *serpent* promoter (Fig S4a-b). Interestingly, loss of N-Cadherin in hemocytes did not affect the migration parameters significantly with hemocytes migrating at similar average speeds and persistence with similar fractions of cells displaying positive mean 3D correlation (Fig 4a-b Fig S4c). However, there was a significant change in the organization of cells within the cluster by stage 11; in the control hemocytes display a preferential distribution along the main axis of migration (Fig 4c); however, upon reduction in N-Cadherin levels, this preferential distribution was lost (Fig 4c,e,f, Video 8) and the hemocyte cluster did not achieve the elongated shape by early stage 11 unlike in the control embryos (Fig 4g). Since N-Cadherin junctional levels are tightly temporally restricted to the elongation phase, we asked what would happen if N-Cadherin levels were increased specifically in the hemocytes (Fig S4a-b)^32^. In hemocytes where N-Cadherin levels were higher than in the control, the cells migrated marginally faster, but with lower persistence (Fig 4a,b). The frequency of cells migrating with higher spatial correlation remained the same when N-Cadherin levels were upregulated (Fig S4c). Interestingly, the cells again dispersed across all quadrants and the cluster did not elongate significantly suggesting that optimal levels of adhesion between the hemocytes is essential for persistent cell movements (Fig 4d-g, Video 8).

**Figure 4.**
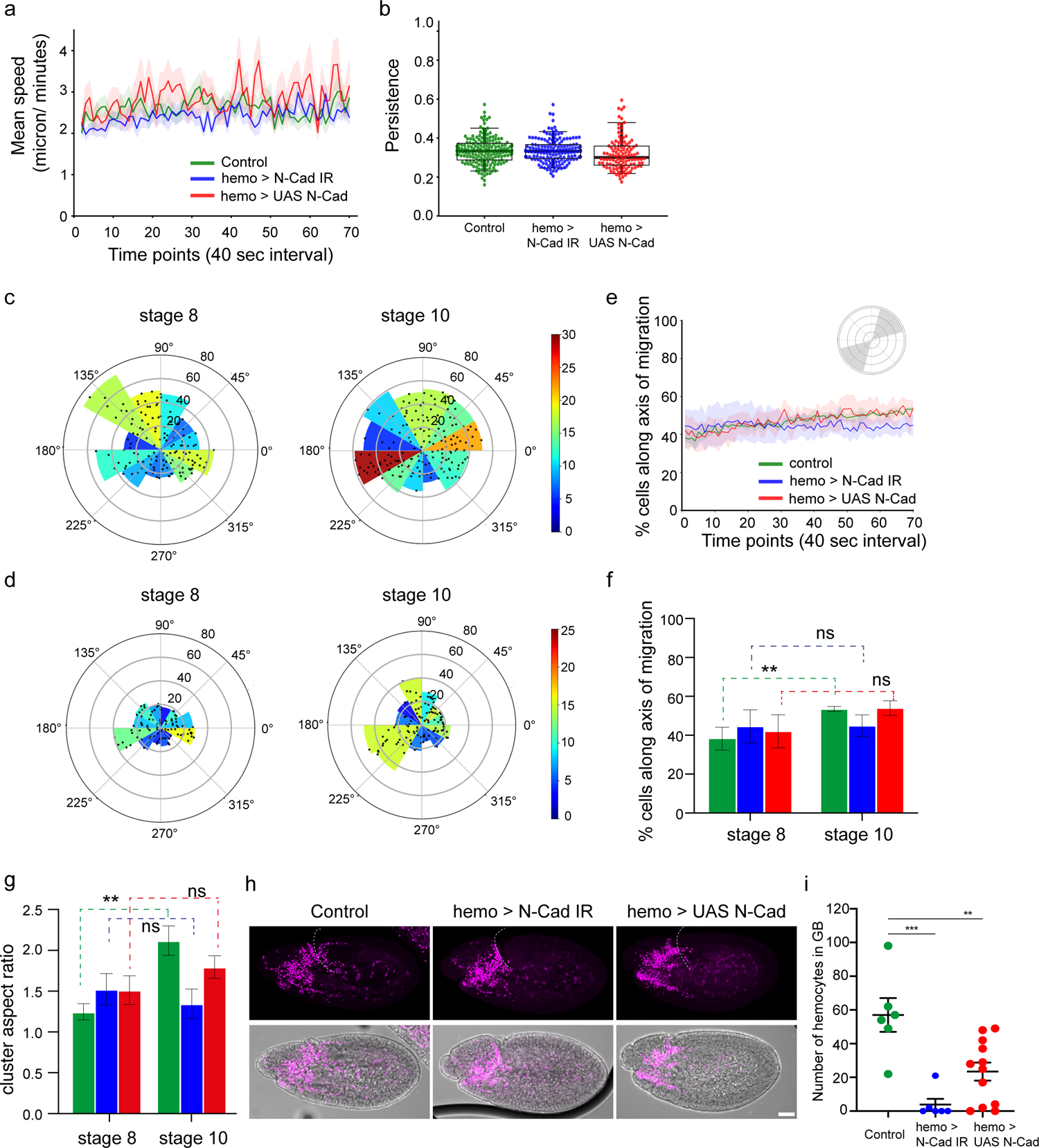
**(a, b)** Line graph showing the mean speed **(a)** and persistence **(b)** of cells between *srpHemo>Valium (green), srpHemo>NCAD RNAi (blue),* and *srpHemo>N-CAD (red)* embryos over time from from stage 8 (0 minutes) and stage 10 (46 minutes). The shaded area around the lines represents the standard error of the mean (SEM). n=3 embryos for *UAS RNAi* and 2 embryos for *UAS N-Cad*. **(c,d)** Polar histogram illustrating the spatial distribution of nuclei between *srpHemo> NCAD RNAi* **(c)**, and *srpHemo>N-CAD* **(d)** at time 0 minutes (stage 8) and 46 minutes (stage 10). Each dot represents the centroids of nuclei segments. The radial axis indicates the physical distance of the displacement in microns, and the angular axis represents the direction in degrees. Colour scale on the right indicates the nucleus number in each 30-degree angle (histogram bin=12). **(e)** Line graph showing the percentage of cells between *srpHemo>Valium (green), srpHemo>NCAD RNAi (blue),* and *srpHemo>N-CAD (red)* embryos in the range of polar histogram from 15-75 degree (1st quadrant, grey shade area in cartoon) and 195-255 degree (3rd quadrant, grey shade area in cartoon) over time from stage 8 (0 minutes) and stage 10 (46 minutes). The shaded area around the line represents the standard error of the mean (SEM). **(f)** Bar graph comparing the percentage of cells in **(e)** at two specific time points, initial (t0) and final (70 minutes). Error bars denote the standard error of the mean (SEM). Statistical significance between stage 8 (0 minutes) and stage 10 (46 minutes) is indicated with ** (p < 0.01). n = 3 embryos. **(g)** Quantification of the shape of the hemocyte cluster in stage 8 and 10 from live imaging of *control* (green), *srpHemo >NCAD RNAi* (blue), and *srpHemo>N-CAD* (red) embryos. n = 3 embryos. **(h,i)** Confocal microscopy images of z projections of fixed lateral stage 12 embryos, from the control (con), *srpHemo>NCAD RNAi,* and *srpHemo>N-CAD* embryos. Macrophages are labeled in red by the expression of srpHemo:3xmCherry. The white dotted line indicates the edge of the germband. Quantification on the right **(i)** shows that both reduction and over expression of N-Cadherin results in reduced macrophage entry into the germband. n = 8-12 embryos.

To understand whether alterations in N-Cadherin levels would affect how the hemocytes are distributed in the embryo, we assessed hemocyte numbers in the germband at the end of stage 11 just before the germband starts to retract. Interestingly, there was a very strong decrease in the number of cells found within the germband in both cases suggesting that optimal levels of N-Cadherin are essential for efficient dispersal of hemocytes along its presumptive route (Fig 4 h,i). Loss of N-Cadherin did not affect overall hemocyte numbers assessed in fixed stage 13 embryos, hence it is unlikely that N-Cadherin has a role in hemocyte specification, proliferation or cell death (Fig S4d).

Our data shows for the first time that *Drosophila* embryonic hemocytes move and elongate as a cluster, behaving more epithelial initially and then transitioning to a mesenchymal phenotype. As a cluster, they are packed densely, display intercalation behaviour commonly seen in epithelial sheets undergoing convergent extension and classical Adherens junctions compartments whose junctional expression is highly dynamic. As the cells leave the cluster in chains, the junctional complexes are dismantled, and the frontal cells acquire dynamic protrusions. The cell cluster displays mechanically distinct behaviours including neighbour exchanges through junctional shrinkage, cell divisions and highly aligned movements in the direction of the cluster movement. Some aspects of hemocyte movement described here are highly reminiscent of neural crest migration and EMT which has been very well studied^33–36^ suggesting that a common toolbox of mechanical and signalling pathways could underlie both these morphogenetic processes. We have not observed planar polarised localization of N-Cadherin in this cluster in our fixed embryos, however since the junction shrinkage that we observe happen extremely quickly, it is possible that N-Cadherin and Myosin polarize for brief periods of time enabling intercalation. While we have not seen a change in the migration parameters of the cluster as a whole when we interfere with N-Cadherin levels, it is possible that N-Cadherin has a role in migration within a specific subset of cells or during discrete periods, generating and transducing forces leading to cluster elongation.

We have not addressed the possible upstream regulators driving this process and studies from systems such as neural crest cells suggest a plethora of both mechanical and chemical signals which could be involved^24,25,27^. Interestingly, cluster elongation seems to orient the bulk of hemocytes along the developmentally hardwired routes of migration. While previous studies have shown a role for Pvfs in guiding hemocytes along their migration routes^8,23^, our study suggests that initial delamination and hemocyte positioning could be a mechanically distinct phenomena which has more in common with epithelial sheets than single cells migrating towards chemotactic cues. However, it is possible that it is an interplay between the strength of adhesive interactions and chemical signalling that dictates the mode of hemocyte migration.

## Acknowledgements

We thank PF. Lenne (IBDM, France) and D. Siekhaus (UCLA, USA) for reagents, and C. Schwayer, M. Smutny and D. Siekhaus and members of our lab for comments on the manuscript. We obtained antibodies from the Developmental Studies Hybridoma Bank (The University of Iowa, USA) created by the NICHD of the NIH. Stocks were obtained from the Bloomington Drosophila Stock Center (NIH P40OD018537) and the Vienna Drosophila Resource Center. Information provided by FlyBase has been crucial to the work. We are grateful to the CAMDU (Computing and Advanced Microscopy Unit) at Warwick Medical School for their support and assistance in this work. This work was supported by the Biotechnology and Biological Sciences Research Council (BBSRC) MIB Doctoral Training Partnership (BB/T00746X/1) to J.S.A, and BBSRC research grant awards (BB/T018070/1 and BB/W017482/1) to A.R., Engineering and Physical Sciences Research Council research grants (EP/V043498/1 and EP/Y002245/1) to D.K., and the Warwick Quantitative Biomedicine Programme funded by the Wellcome Trust Institutional Strategic Support Fund (ISSF) to A.R and D.K.

## Author Contributions

A.R. conceived the project and A.R, J.H and H.S designed the research. A.R, J.H and H.S performed the experiments and analysed the data. X.M, D.V.K and J.S.A performed computational image processing and analysis. A.R wrote the manuscript with input from D.V.K. All authors edited the manuscript.

## Declaration of Interest

The authors declare no competing interests.

## Supplemental Data

### Supplemental Figures

**Figure S1.**
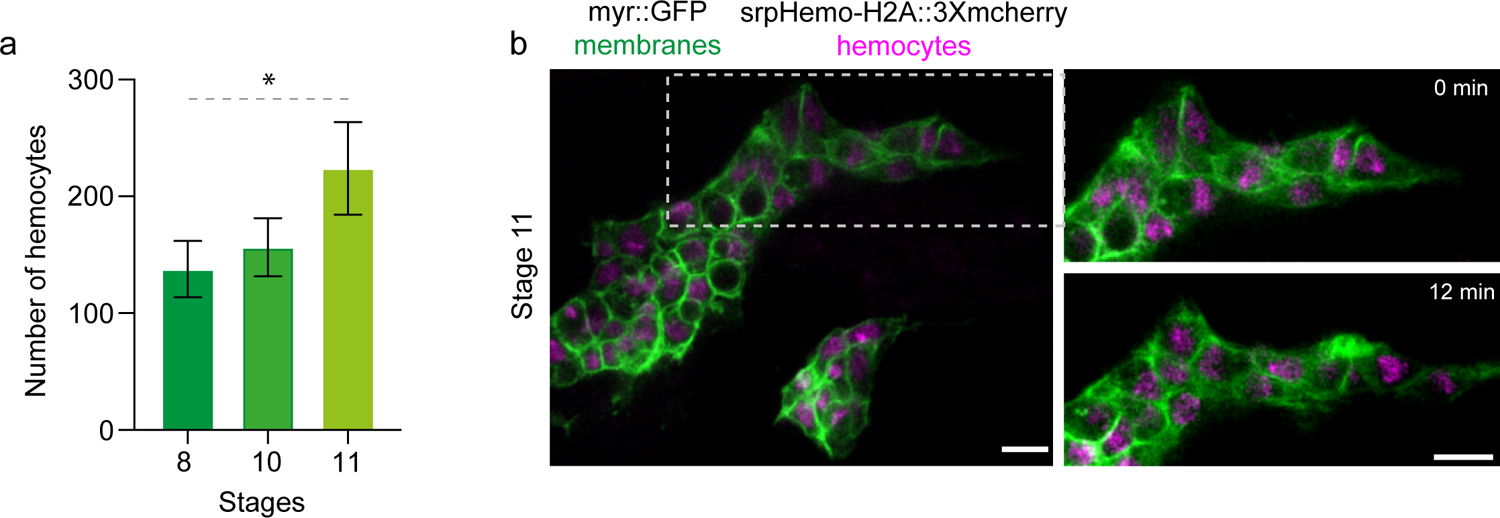
**(a)** Quantification of total number of hemocytes in *srpHemo>myr::GFP; srpHemo-H2A::3xmCherry* embryos across stages 8,10 and 12. n= 4-6 embryos. **(b)** Representative images from multiphoton time-lapse imaging of a *srpHemo>myrGFP; srpHemo-H2A::3xmCherry* embryo from 0 min to 12 min showing hemocytes migrating out of the cluster in chains. Scale bars = 10 µm.

**Figure S2.**
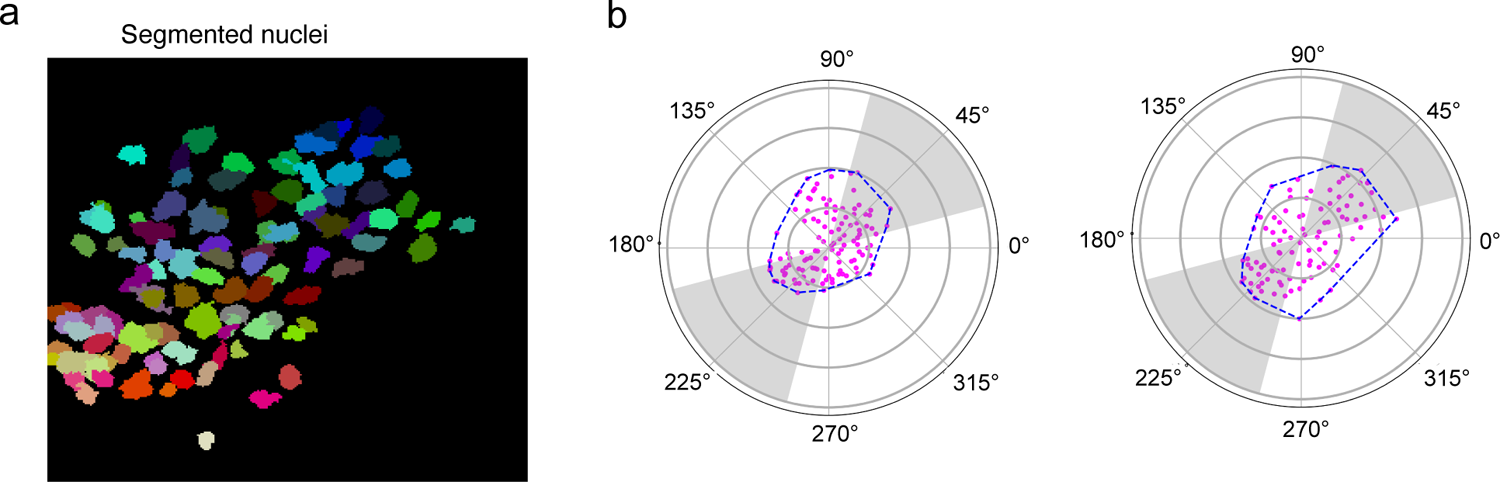
**(a)** Instance segmentation z projection of nuclei predicted by pretrained 3D U-Net model. Each distinct coloured segment represents a nucleus identified by the model. **(b)** Polar distance illustrating the spatial distribution of nuclei between stage 8 (0 mintes) and stage 10 (46 minutes). from live imaging of *srpHemo-H2A::3XmCherry* embryos. Each magenta dot represents the centroids of nuclei segments. The radial axis indicates the physical distance of the displacement in microns, and the angular axis represents the direction in degrees. The blue polygon outlines the collective area encompassing the population of nuclei, illustrating the general shape and displacement of the nuclei distribution. The percentage of cells was calculated in specific grey shade areas in the 1st quadrants (15-75 degree) and 3rd quadrants (195-255 degree) over consecutive time points.

**Figure S3.**
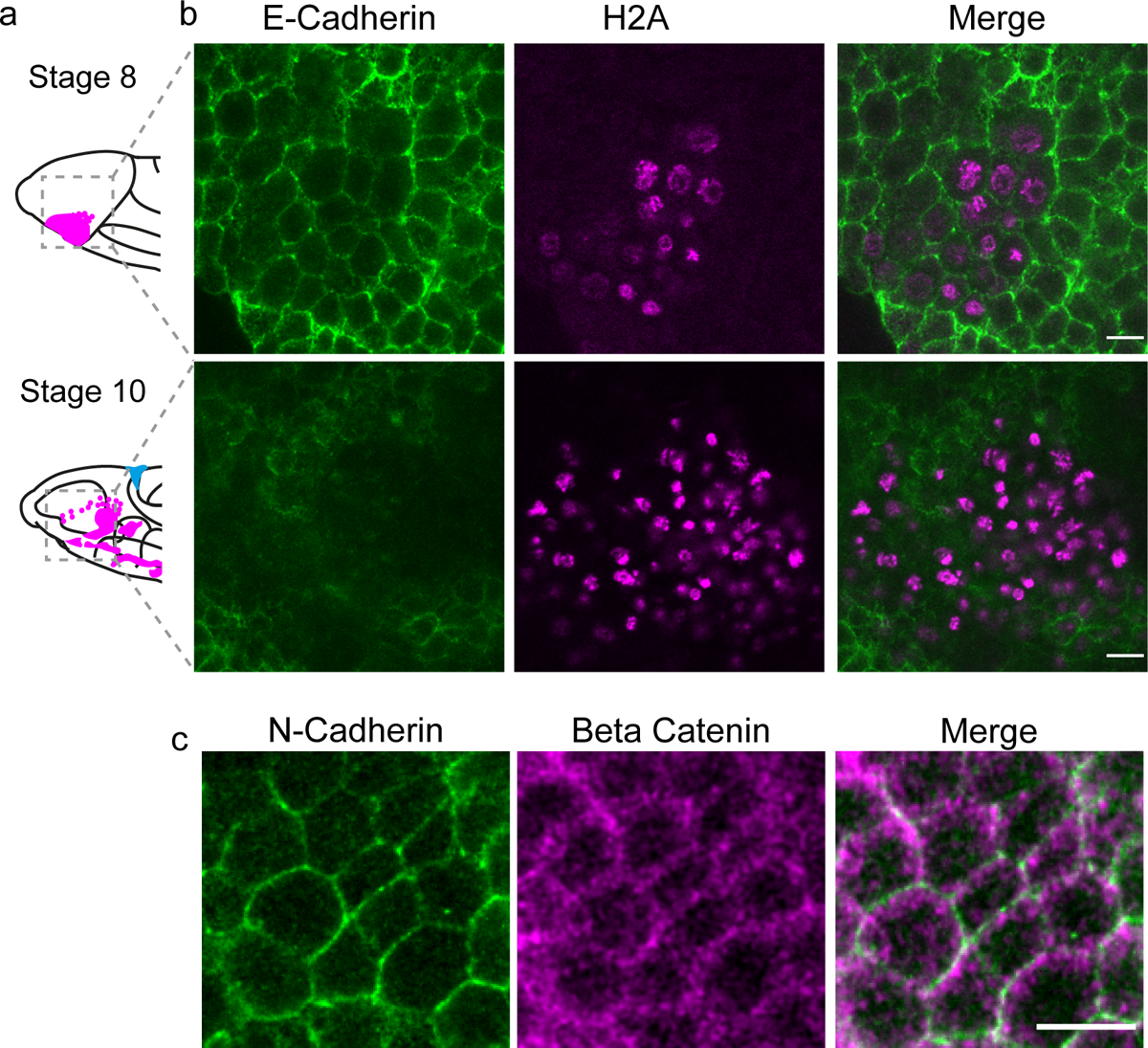
**(a)** Cartoon showing the shape of the hemocyte cluster at stage 8 and 10 (Magenta). Dotted boxes indicate the regions of interest shown in (a). **(b)** Representative image of a single z slice of fixed confocal microscopy images of *E-Cadherin::GFP; srpHemo-H2A::3XmCherry* embryos at stage 8 and 10. *srpHemo-H2A::3XmCherry* labelled hemocyte nuclei (magenta). E-Cadherin localised to hemocyte membranes at stage 8 (green), but is absent by stage 10, with both channels merged. Scale bars = 10 µm. **(c)** Single z slice of fixed confocal microscopy image of a stage 8 *N-Cadherin::GFP* (Green) embryo within the hemocyte cluster immunostained for Armadillo (Beta-Catenin) (Magenta), along with an image with both channels merged. Scale bars = 10 µm.

**Figure S4.**
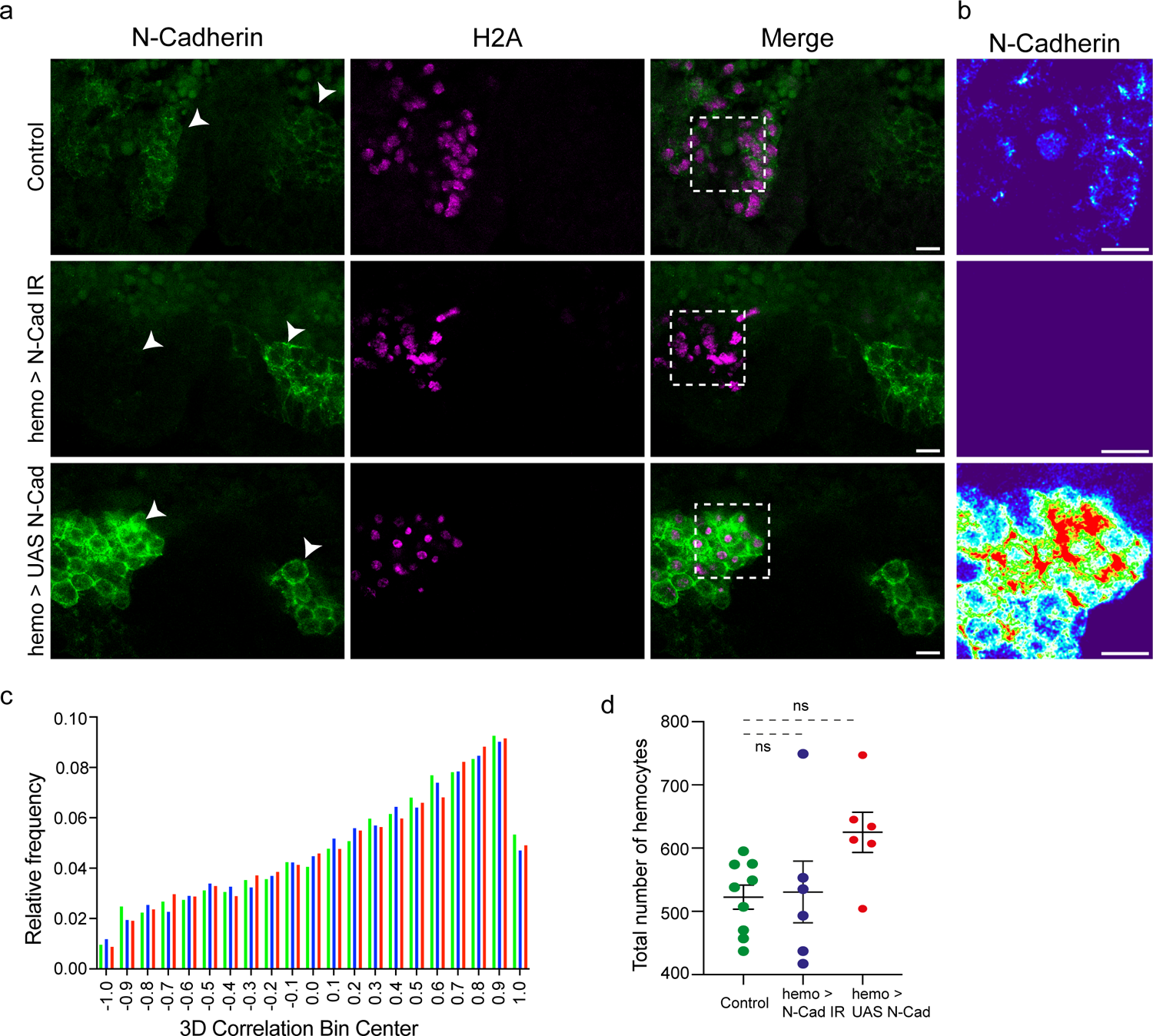
**(a)** Representative z-projections of N-Cadherin staining (Green) in stage 8-9 *srpHemo-GAL4>Valium, srpHemo-GAL4>NCAD RNAi,* and *srpHemo-GAL4>NCAD* embryos. All genotypes carried *srpHemo-H2A::3XmCherry*, labelling hemocyte nuclei (Magenta). White arrows indicate the hemocyte cluster (left) and the mesoderm (right). Scale bars = 10 µm. **(b)** Heatmap of N-Cadherin::GFP fluorescence intensity from a single z slice within the hemocyte cluster in a region of interest indicated by the white dotted line boxes in the merged images of (A). Scale bars = 10 µm **(c)** Relative frequency of 3D velocity cluster correlation from individual cells over the entire 46 minutes from *srpHemo-GAL4>Valium* (green), *srpHemo-GAL4>NCAD RNAi* (blue), and *srpHemo-GAL4> UAS-N-CAD* (red) embryo **(d)** Quantification showing that reduction and over expression of N-Cadherin does not change total hemocyte numbers in the *srpHemo-GAL4>NCAD RNAi,* and *srpHemo-GAL4>NCAD* embryos compared to *srpHemo-GAL4>Valium* controls.

**Video 1:** Multiphoton time-lapse imaging of a *srpHemo>myrGFP; srpHemo-H2A::3xmCherry* embryo from stage 8 (0 mins) to late stage 11 (72 mins), in which *GFP* labels hemocyte cell membranes (green) and *mCherry* labels hemocyte nuclei (magenta). Frame rate= 40 seconds, scale bars = 10 µm.

**Video 2:** Multiphoton time-lapse imaging of control *srpHemo-H2A::3xmCherry* embryo stage 8 to stage 10, Frame rate= 40 seconds, scale bars = 10 µm.

**Video 3:** Multiphoton time-lapse imaging of control *srpHemo-H2A::3xmCherry* embryo showing cell division events. Frame rate= 40 seconds, scale bars = 10 µm.

**Video 4:** Multiphoton time-lapse imaging of a *srpHemo>myrGFP; srpHemo-H2A::3xmCherry* embryo showing T1 transitions indicating junction shrinkage and cell rearrangements (magenta asterisk and lines). Frame rate= 40 seconds, scale bar=5 microns.

**Video 5:** Multiphoton time-lapse imaging of an *N-Cadherin::GFP; srpHemo-H2A::3XmCherry* embryo. Frame rate= 40 seconds, scale bar=5 microns.

## Materials and Methods

### Fly strains and husbandry

Flies were reared at 25°C at 65% humidity on standard cornmeal-based food. Embryos were collected on standard plates containing apple juice, agar, sugar, and Nipagen, with inactive yeast applied to the surface of the plates. Embryo collections were conducted at 29°C with 3-hour collections. The fly lines used here are described below: *srpHemo-GAL4*^23^, *srpHemo-H2A::3xmCherry* ^37^and *DE-Cadherin::GFP* ^38^was provided by D. Siekhaus. *NCAD::eGFP* and *UAS-NCAD* was provided by P-F. Lenne^32^. The following lines were obtained from the Bloomington Stock Centre: *UAS-N-Cadherin* RNAi (TRiP.HMS02380)(BL 41982), Valium-10 control (BL 35786), *UAS-myr::GFP* (BL 32197), and *UAS-moe::GFP* (BL 31776).

### Immunofluorescence

Embryos were dechorionated in 50% bleach for 5 minutes and were subsequently washed with water. For most antibody stainings embryos were fixed via a 45-minute incubation in a 5% methanol-free formaldehyde and 50% heptane mixture followed by an ethanol devitellinization as described previously^39^. Embryos for Phalloidin, *E-Cadherin::GFP* and *N-Cadherin::GFP* stainings were devitellinised by hand. For anti-Armadillo stainings, embryos were heat fixed, as previously described^40^ by incubating dechorionated embryos in 10ml of boiling Heat fixation buffer (68 mM NaCl, 0.03% Triton-X) for 10 seconds before cooling to ice cold. The buffer was replaced by 5ml of Heptane, and the embryos were devitellinised via the addition of 10ml of methanol and vigorous shaking for 1 minute. Fixed embryos which had completed germband extension were staged for imaging based on the invagination of the stomodeum as well as germband retraction away from the anterior as described previously^39^.

The primary antibodies used were: Rabbit anti-GFP (Invitrogen A-11122, 1:300), Rat anti-DN-Cadherin (DSHB DN-Ex #8, 1:10), Mouse anti-Armadillo (DSHB N2-7A1, 1:75). The secondary antibodies used were: Goat anti-Rabbit 488 (Invitrogen A-11008, 1:500), Goat anti-Mouse 633 (Invitrogen A-21050, 1:100), and Goat anti-Rat 405 (Invitrogen A-48261, 1:250). Phalloidin 405 (Invitrogen A-30104) was utilized at a dilution of 1:250. Embryos were mounted in Vectashield Mounting Medium (Vector Labs, Burlingame, USA). Images were acquired with an Olympus FV3000 inverted confocal microscope, with a 20X/0.5 air objective or a 60X/1.35 oil immersion objective as required.

### Time-Lapse Imaging

Embryos were dechorionated in 50% bleach for 5 minutes and were subsequently washed with water and were then mounted onto a coverslip lateral side up and were covered in halocarbon oil 700 (Sigma-Aldrich). The embryos were imaged on an Olympus inverted multiphoton microscope with an Olympus 60X/1.42 oil immersion objective at 28°C with a step size of 0.5 µm. GFP and mCherry were imaged at 960 nm and 1140 nm excitation wavelengths respectively, with a frame rate of 40 seconds for *N-Cadherin::GFP* and srpHemo-H2A::3xmCherry time-lapses and at 50 seconds for *srpHemo>GMA* and *srpHemo>myrGFP* time-lapses. Excitation intensities were adjusted for tissue depth.

### Image Process and Analysis of Macrophage Migration

Time-lapses of embryos in which the macrophage nuclei were labelled with *srpHemo-H2A::3XmCherry* were typically acquired as 169.71 x 169.71 x 31μm^3^ 3D-stacks with a constant 0.33 x 0.33 x 1μm^3^ voxel size with a frame rate of 40 seconds for approximately 1 hour. Images acquired from multiphoton microscopy were further processed using Imaris software (Bitplane) to visualize the recorded channels in 3D. Briefly, the time-lapse of an embryo was rotated and aligned along the AP axis for tracking analysis and movies were cropped in time to a period of typically 55 minutes from the original movie to calculate migration parameters.

### Image 3D Segmentation and Analysis

The 3DeeCellTracker program was used to perform 3D image segmentation of time-lapse microscopy data, facilitating cell division and migration tracking in a dynamic cell environment ^41^. Prior to segmentation, the nuclear channel of the 3D image stack was manually annotated to create a robust training dataset. Annotations were performed on two volumes for each image stack, one for the training dataset and the other for the validation dataset, which were selected to represent the diversity of nucleus morphology. The annotation image sets were then used to train the 3DeeCellTracker model. The training process utilized the 3D-UNet, a specialized convolutional neural network (CNN), designed for the segmentation of 3D volumetric data. Training was conducted iteratively and adjusted dynamically based on the validation loss observed during the training epochs. The pretrained model was used to the prediction of intensity-based segmentation on entire image stacks. The training and prediction utilized the full capabilities of a workstation equipped with an RTX A5000 graphics card (NVIDIA), featuring 24 GB of memory. In some areas, segmentation was corrected manually in the final binary images, and nuclei that appear to be in the direction of the ventral nerve cord were removed manually from the segment image for further analysis.

The resulting binary image stacks were analysed using a custom Python script to obtain the centroids of the nuclei positions in XYZ dimensions. The 2D density heat map was then generated based on the XY position of the nucleus using Gaussian kernel density estimation (KDE) to calculate the probability density of the point characteristics around each other between consecutive time points. To generate the polar histogram and the polar distance map, we determined the median of nuclei XY position as the centre of a Cartesian coordinate system and calculated the distance and the radian between the centroids of nuclei in each quadrant and the centre for each time frame respectively. To observe and quantify cell movements, the Delaunay triangulation was calculated using the SciPy library spatial module, where the centroids of nuclei function as vertices.

### Cell Tracking and Analysis

Macrophage nuclei were extracted using the spot detection function and tracks generated manually in 3D over time. Due to limitations in imaging parameters, not all macrophages could be tracked. Tracks of macrophages which migrate towards the dorsal vessel, ventral nerve cord and to the anterior of the head were omitted. In all cases *H2A::3xmCherry* labelling used to follow individual cells over time, and nuclei positions in XYZ-dimensions were generated by the Imaris tracking tool.

Cell speeds, persistence, and velocity correlation were calculated using custom Python scripts. Briefly, instantaneous velocities from single cell trajectories were averaged to obtain a mean instantaneous velocity value over the course of measurement. To calculate persistence values, single cell trajectories were split into segments of equal length (10 frames) and calculated via a sliding window as the ratio of the distance between the macrophage start-to-end distance over the entire summed distance covered by the macrophage between each successive frame in a segment^39^. Obtained persistence values were averaged over all segments in a single trajectory. This analysis yielded values between [0,1], with higher movement directionality closer to 1. To investigate the collective movement and potential mechanical interaction between cells, we calculated the velocity correlation coefficient between the direction of movement of each individual cell and the average movement direction of the collective as described previously ^42^. This coefficient value of +1 indicates a perfect correlation (cells moving in the same direction), 0 indicates no correlation, and −1 indicates a perfect anti-correlation (cells moving in the opposite direction) in any time frame.

### Measurement of Junctional Fluorescence Intensities

*N-Cadherin::GFP; srpHemo-H2A::3XmCherry* embryos were imaged as 3D-stacks with a constant voxel size of 0.207 x 0.207 x 0.5 µm^3^. Each 3D stack was rotated to align with the anterior-posterior axis. Hemocytes were identified by *srpHemo-H2A::3XmCherry* nuclei labelling. Maximum projections were produced from 3-10 slices to capture a single layer of cells.

### ROI analysis of N-Cadherin::eGFP fluorescence intensity

Single cell-layer maximum projection images were used to quantify *N-Cadherin::eGFP* fluorescence intensities by tracing an ROI around a hemocyte-hemocyte cell contact and measuring Mean Gray Value in ImageJ/Fiji. *N-Cadherin::eGFP* fluorescence intensity was quantified at non-hemocyte mesodermal cell contacts from the same projection as hemocyte contacts. *N-Cadherin::eGFP* fluorescence intensity at a given hemocyte contact was normalised to the mean intensity of N-Cadherin in mesodermal contacts at the same depth.

### Measurement of junctional intensity profiles

The junctional fluorescence intensity profiles of *N-Cadherin::eGFP* at hemocyte-hemocyte cell contacts was quantified using linescan analysis in ImageJ/Fiji in single-cell layer maximum projections of *N-Cadherin::eGFP* embryos as previously described ^43^with line length set to 30 pixels (6.21 µm).

### Cluster analysis

Cluster analyses were performed in ImageJ/Fiji using the measurement tool to measure the length and width of each cluster. Measurements were taken from the nuclear channel in 3D segmented images to calculate aspect ratio (length of cluster along the axis of migration/ width of cluster perpendicular to the axis of migration). Numbers of macrophages in each group was calculated using the spot function, Imaris software (Bitplane).

### Statistical Analysis

Statistical tests as well as the number of embryos/ cells assessed are listed in the Figure legends. All statistical analyses were performed using GraphPad Prism. The experiments were not randomized and no statistical analysis was done to predetermine sample size. Data points from individual experiments / embryos were pooled to estimate mean and s.e.m. Error bars in all graphs represent the standard error of the mean. Unpaired t-test was used to calculate the significance in differences between two groups and One-Way ANOVA followed by Dunnett post tests were used for multiple comparisons.

## Notes

### Competing Interest Statement

The authors have declared no competing interest.

